# Recognition of the centromere-binding protein Ndc10 by Scm3^HJURP^ directs Cse4^CENP-A^ deposition at native centromeres

**DOI:** 10.64898/2026.10.30.763002

**Authors:** Jiaxi Cai, Ziyi Wang, Lily Cheng, Tianjian Xu, Cindy Zhang, Stephen M. Hinshaw, Raymond T. Suhandynata, Arshad Desai, Huilin Zhou

## Abstract

Kinetochores connect centromeres to spindle microtubules, ensuring accurate chromosome segregation. Kinetochore assembly at centromeres depends on deposition of CENP-A, a histone H3 variant. Despite considerable progress, how CENP-A is specifically assembled on native centromeres remains an open question. Centromere-specific assembly of CENP-A occurs spontaneously in a cell-free extract system from the yeast *Saccharomyces cerevisiae*, but with low efficiency, suggesting that one or more factors are limiting. Here, we show Scm3^HJURP^, a specific chaperone for the centromeric histone Cse4^CENP-A^, is one such limiting component. The addition of a complex of Scm3, Cse4 and histone H4 strongly stimulates Cse4 assembly in a centromeric DNA sequence- and dose-dependent manner; in addition, improvements in both Cse4 and kinetochore assembly were more prominent in mitotic versus interphase (G1) extracts. Using the complemented cell-free reconstitution system, we show that distinct Scm3 interfaces with Cse4 and with the Ndc10 subunit of the yeast centromere DNA sequence-recognizing CBF3 complex are essential for centromere-specific Cse4 assembly and cell viability. Thus, recognition of Ndc10 directs Scm3-dependent and locus-specific centromeric histone deposition.

**SIGNIFICANCE STATEMENT:** Accurate chromosome segregation depends on the assembly of kinetochores at centromeres. In budding yeast, centromeres are specified by DNA sequence, but how sequence recognition is coupled to deposition of the centromeric histone Cse4 remains incompletely understood. Using a cell-free assembly system, we show that Scm3 promotes centromeric Cse4 assembly through distinct regions that bind Cse4/H4 and the centromere-binding protein Ndc10. Mutations that disrupt Ndc10 binding impair Cse4 assembly in extracts, reduce Scm3 association with centromeres in cells, and compromise viability. These findings support a mechanism in which a histone chaperone engages a sequence-specific centromere-binding complex to direct centromeric histone assembly.

## INTRODUCTION

Eukaryotic centromeres are specialized chromatin regions that direct kinetochore assembly. Kinetochores form the physical linkage between spindle microtubules and sister chromatids during chromosome segregation (1, 2). Kinetochores are organized into inner and outer layers, where the inner kinetochore assembles on centromeric DNA, and the outer kinetochore mediates microtubule attachment. During each cell cycle, kinetochores in yeast disassemble to accommodate centromeric DNA replication in S phase and subsequently reassemble on newly replicated centromeres in preparation for mitosis (3). While the composition and function of kinetochores have been extensively characterized, the mechanism by which inner kinetochore proteins assemble on the native centromeric DNA and not elsewhere on the chromosome is not fully resolved. Furthermore, how inner kinetochore assembly accommodates the cell cycle so that premature stabilization does not block DNA replication has not been fully understood. To address these knowledge gaps, it is critical to study the initial event of kinetochore assembly on the centromeric DNA in a native environment.

Eukaryotic centromere identity is epigenetically specified by CENP-A, a histone H3 variant that forms a heterodimer with histone H4 within the centromeric nucleosome (4). The CENP-A chromatin provides a platform for recruiting inner kinetochore components that are broadly conserved across eukaryotes (1, 2, 5), including CENP-C (6), and the multi-subunit constitutively centromere-associated network (CCAN) (7). Homologous components are present in the budding yeast *Saccharomyces cerevisiae*, including Cse4^CENP-A^ (8), Mif2^CENP-C^ (9), and the Ctf19c^CCAN^ complex (10–12). Although these components, their organization, and their structures have been well characterized, this knowledge has yet to translate into a stepwise mechanistic understanding of how these inner kinetochore components are assembled on native centromeric DNA.

A defining feature of the inner kinetochore is its multivalency, arising from extensive protein-protein and protein-DNA interactions. Structural and biochemical studies have provided insights into the organization of subcomplexes within the kinetochore, including interactions between the N-terminus of Cse4 and the Ame1/Okp1 subcomplex (13), as well as the association of Mif2 with Cse4 (14, 15). In yeast, the connection between Mif2 and Ctf19c/CCAN may be mediated by Chl4-Iml3 and Ame1-Okp1 (16, 17), whereas human CENP-C interacts with the four-subunit CENP-HIKM subcomplex (18). Moreover, many inner kinetochore proteins exhibit DNA-binding activity (14, 19–22). Together, these observations support a model in which multivalent protein-protein and protein-DNA interactions stabilize the inner kinetochore and contribute to its centromere specificity. However, establishing how these interactions contribute to cell cycle-appropriate kinetochore assembly requires the reconstitution of kinetochores on native centromeric DNA in a system which recapitulates endogenous regulation while providing experimental control.

Alongside progress in characterizing the inner kinetochore subcomplexes (10, 11, 23), CENP-A nucleosomes have been reconstituted on native centromeric DNA from purified components (24, 25). These particles are unstable, however: Cse4 nucleosomes assembled on CEN3 DNA show no assembly preference over strong positioning sequences and dissociate far more rapidly (24), and a cryo-EM structure on native CEN3 required a stabilizing antibody fragment to survive sample preparation (25). These reconstitutions revealed that the AT-rich character of centromeric DNA, a feature broadly conserved across eukaryotes (23, 26, 27), is unfavorable for stable nucleosome wrapping. This property has contributed to a long-standing debate about the composition and architecture of CENP-A chromatin in vivo (24, 26, 28–32). Because assembly of CENP-A on centromeric DNA is required for kinetochore formation, and because purified systems lack the cellular factors that act on this step, an approach is needed that permits CENP-A assembly on native centromeric DNA to be studied in the presence of such factors.

The point centromere in budding yeast offers an ideal system for such an approach, as it has well-defined DNA sequences (33). Building on this, kinetochore proteins, including Cse4^CENP-A^, have been shown to assemble on centromeric DNA in whole-cell extracts (WCEs), which recapitulated the physiological features of kinetochores, including centromeric DNA sequence specificity, cell-cycle regulation, and microtubule-binding activity (34–37). This approach has since been extended to single-molecule imaging, which resolved the arrival order of individual components on CEN DNA and showed that Cse4 occupancy depends on the chaperone Scm3 and is stabilized by centromere-bound inner kinetochore proteins (38, 39). Moreover, our recent stoichiometric analysis showed that most if not all assembled kinetochore complexes are incomplete, showing a relatively low efficiency in the assembly of Cse4^CENP-A^ on the centromeric DNA (40). Thus, the availability or activity of one or more factors appears to limit efficient kinetochore assembly in this cell-free system, highlighting the need to identify these factors and characterize their roles in centromeric DNA-dependent assembly to better understand the underlying molecular mechanisms.

The budding yeast centromere consists of three conserved elements (CDEI-III). CDEI binds Cbf1, CDEIII binds the sequence-specific CBF3 complex, and the AT-rich CDEII element is occupied by a single Cse4 nucleosome (33, 41–44). The CBF3 complex is essential for cell viability, and it comprises four subunits (Ndc10, Cep3, Skp1 and Ctf13). CBF3 directly contacts a conserved CCG motif in the CDEIII region to specify the native centromeric DNA, a physiological requirement for the assembly of Cse4 and the rest of the kinetochore. Additionally, several proteins have been shown to affect kinetochore assembly in vivo. For instance, Sgt1 has been implicated in CBF3 regulation and inner kinetochore assembly (45, 46), while Scm3, a Cse4-specific chaperone and functional ortholog of metazoan HJURP, is required for centromeric Cse4^CENP-A^ deposition in vivo (26, 47, 48). Biochemical studies showed that Scm3^HJURP^ forms a complex with Cse4^CENP-A^ and histone H4 (31, 49), and this interaction is mutually exclusive with histone H2A/H2B (26), suggesting that Scm3 may deliver Cse4/H4 to form the Cse4-specific nucleosome with H2A/H2B (24, 28). Scm3 has also been reported to possess DNA-binding and CBF3-binding activities (27, 50), suggesting an early role in kinetochore assembly. Consistent with such a role, single-molecule imaging in mitotic extracts showed that Scm3 abundance limits Cse4 occupancy at CEN DNA and that Ndc10 usually arrives before Cse4 (38). How Scm3 is directed to centromeric DNA, and which of its interaction surfaces couple centromere recognition to Cse4 deposition, remain undetermined. Here, we use this cell-free system to identify Scm3 as a limiting component for Cse4 assembly on native centromeric DNA. Addition of a recombinant Scm3/Cse4/H4 (SCH) complex strongly stimulates Cse4 deposition in a DNA sequence- and dose-dependent manner. The resulting chromatin supports increased recruitment of both inner and outer kinetochore proteins, with more efficient assembly observed in mitotic compared with interphase (G1) extracts. Using separation-of-function mutants, we find that Scm3 engages Cse4/H4 and the CBF3 subunit Ndc10 through distinct binding interfaces, and that each interface is required for Cse4 assembly in extracts, for Scm3 localization to centromeres in vivo, and for cell viability. Thus, recognition of a sequence-specific DNA-binding factor directs a histone chaperone to deposit CENP-A at a defined chromosomal locus.

## RESULTS

### An Scm3-Cse4-H4 (SCH) complex promotes cell-free assembly of Cse4 on centromeric DNA

To examine kinetochore assembly on native centromeric DNA, we used a cell-free reconstitution assay (34) with bead-immobilized DNA substrates containing wild-type centromere III (CEN), a CDEIII-mutated centromere (MUT), or a non-centromeric autonomously replicating sequence (ARS) control (**Figure 1A**). To determine what protein factors might be limiting for inner kinetochore assembly in this system, we examined the effects of supplementing the whole cell extract (WCE) of cells arrested in mitosis (M-phase) by nocodazole with purified recombinant proteins (**Figure 1B**). Following incubation, DNA bead-associated proteins were recovered and analyzed by immunoblotting to monitor the extent of CEN-specific Cse4 recruitment (**Figure 1C and Supplementary Figure 1**). Addition of Cbf1 or Sgt1 had little detectable effect on the recovery of either Cse4 or Ndc10. In contrast, addition of a Scm3/Cse4/H4 (SCH) complex markedly increased Cse4 recovery from CEN DNA. Recombinant SCH also stimulated Cse4 assembly on MUT or ARS DNA, but to a far lesser degree than on CEN DNA. SCH addition did not appreciably alter Ndc10 recovery, which remained highest on CEN DNA. SCH was next titrated to determine how its concentration affected Cse4 assembly (**Figure 1D** and **1E**). At lower SCH concentrations, increased Cse4 recovery was most prominent on CEN DNA. At the highest concentration tested (600 nM), marked increases in Cse4 recovery were also observed on MUT and ARS DNA, indicating reduced DNA-sequence specificity at elevated SCH concentrations. Because recombinant SCH preparations contained different epitope-tag configurations, their effects on Cse4 assembly were also compared. SCH constructs containing 6xHis-tagged Scm3 and H4 retained preferential Cse4 recovery on CEN DNA and were used for subsequent experiments (**Supplementary Figure 2**).

**Figure 1.**
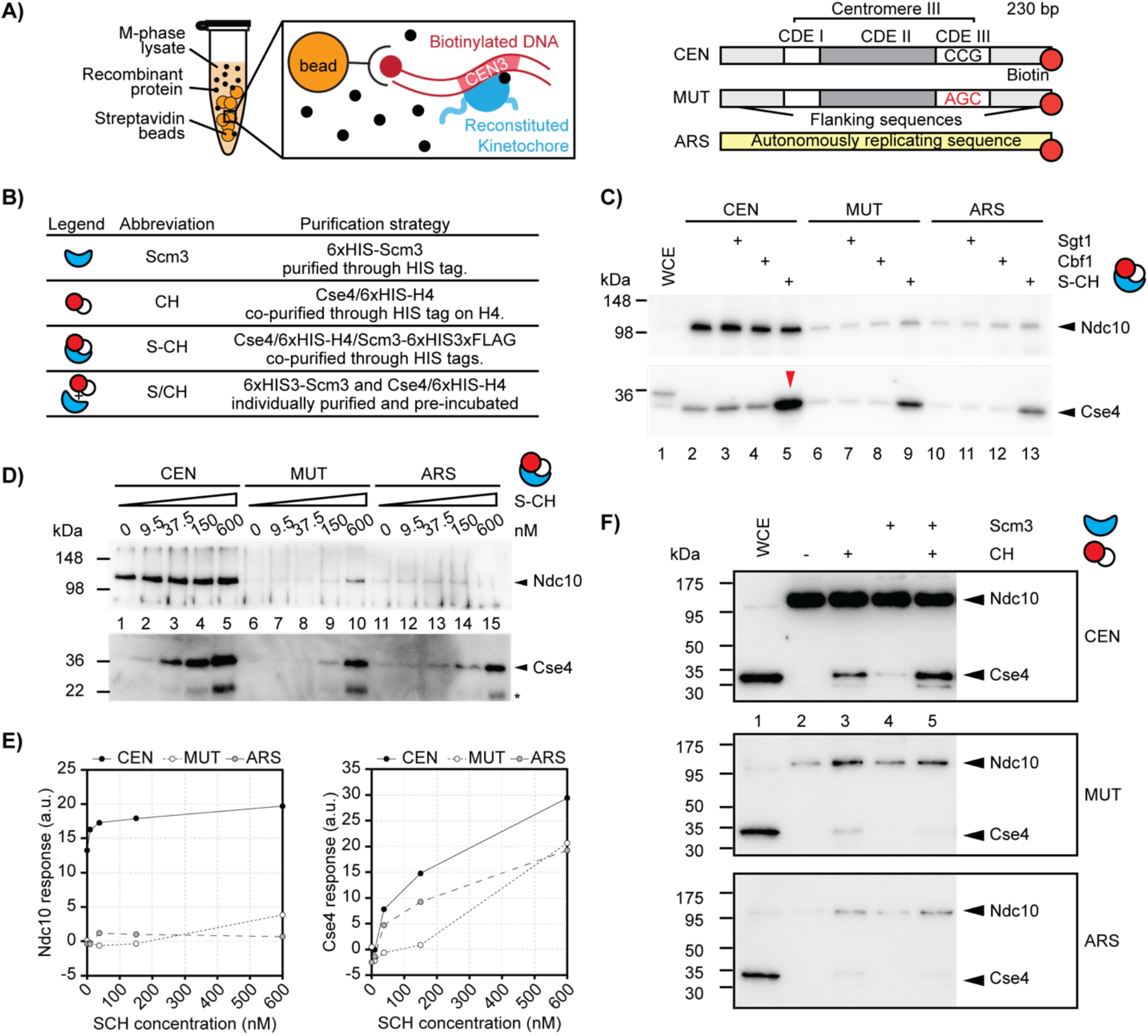
The addition of Scm3/Cse4/H4 (SCH) stimulates the ex vivo reconstitution of the kinetochore. (A) Schematic of the cell-free kinetochore reconstitution assay (left) and DNA substrates (right). Biotinylated, 230-bp DNA substrates containing wild-type centromere III (CEN), centromere III with a CCG-to-AGC substitution in CDEIII (MUT), or a non-centromeric autonomously replicating sequence (ARS) were immobilized on streptavidin-coated beads and incubated with M-phase yeast whole-cell extract (WCE), with or without recombinant proteins. After washing, DNA-associated proteins were released by DNase I digestion and analyzed by immunoblotting. (B) Recombinant protein preparations and symbols used in the reconstitution experiments. Scm3 was purified through an N-terminal 6xHis tag, and Cse4/H4 (CH) was co-purified through 6xHis-tagged H4. S-CH denotes the co-purified Scm3/Cse4/H4 complex, whereas S/CH denotes separately purified Scm3 and CH pre-incubated together before addition to WCE. Epitope-tag configurations are indicated. (C) Effects of recombinant Sgt1, Cbf1, and S-CH on Cse4 assembly. CEN, MUT, and ARS DNA beads were incubated with WCE alone or supplemented with Sgt1 (560 nM), Cbf1 (635 nM), or S-CH (180 nM). DNA-associated Ndc10 and Cse4 were detected by immunoblotting. Lane 1 contains WCE input; lanes 2-5, 6-9, and 10-13 show CEN, MUT, and ARS reactions, respectively. The red arrowhead marks the increased Cse4 signal in the CEN reaction supplemented with S-CH. Ndc10 and Cse4 blots were recorded using different exposure times. (D) Titration of co-purified S-CH. CEN, MUT, and ARS DNA beads were incubated with WCE supplemented with 0, 9.5, 37.5, 150, or 600 nM S-CH. DNA-associated Ndc10 and Cse4 were detected by immunoblotting. Asterisks mark a cross-reactivity/degradation product. (E) Densitometric quantification of Ndc10 (left) and Cse4 (right) immunoblot signals from panel D. Responses are plotted in arbitrary units (a.u.) against S-CH concentration for CEN, MUT, and ARS DNA substrates. (F) Contributions of Scm3 and CH to Cse4 assembly. CEN, MUT, and ARS DNA beads were incubated with WCE alone or supplemented with CH, Scm3, or a pre-incubated mixture of Scm3 and CH. Each recombinant component was added at a final concentration of 100 nM where present. DNA-associated Ndc10 and Cse4 were detected by immunoblotting. Lane 1 contains WCE input; lanes 2-5 show the indicated reconstitution conditions.

The individual contributions of Scm3 and Cse4/H4 (CH) were next examined by adding them individually or together to the reconstitution reaction (**Figure 1F**). Cse4 recovery was minimal in reactions containing no recombinant protein or Scm3 alone, whereas addition of CH produced a modest increase in CEN-associated Cse4. The greatest increase was observed when Scm3 and CH were pre-incubated together before addition to the reaction. Titration of the individual components further showed that Scm3 and CH could be added at concentrations up to 400 nM while largely maintaining preferential Cse4 recovery on CEN DNA relative to MUT and ARS controls (**Supplementary Figure 3**). These results demonstrate that Scm3 and Cse4/H4 cooperate to enhance CEN-specific Cse4 assembly.

### Scm3 and Cse4/H4 stimulate kinetochore assembly in G1- and M-phase extracts

The effect of SCH on kinetochore assembly was next examined using WCEs prepared from cells arrested in G1- or M-phase (**Figure 2A**). SCH addition increased the recovery of Cse4 and Mif2 on CEN DNA in both WCEs. However, CEN-associated Cse4 and Mif2 remained more elevated in M-phase WCEs than in G1 WCEs, regardless of SCH addition. Ctf19 recovery was also assessed by immunoblotting with an anti-Ctf19 antibody, but its detectable signal was near background levels, preventing a reliable assessment of SCH-dependent changes.

**Figure 2.**
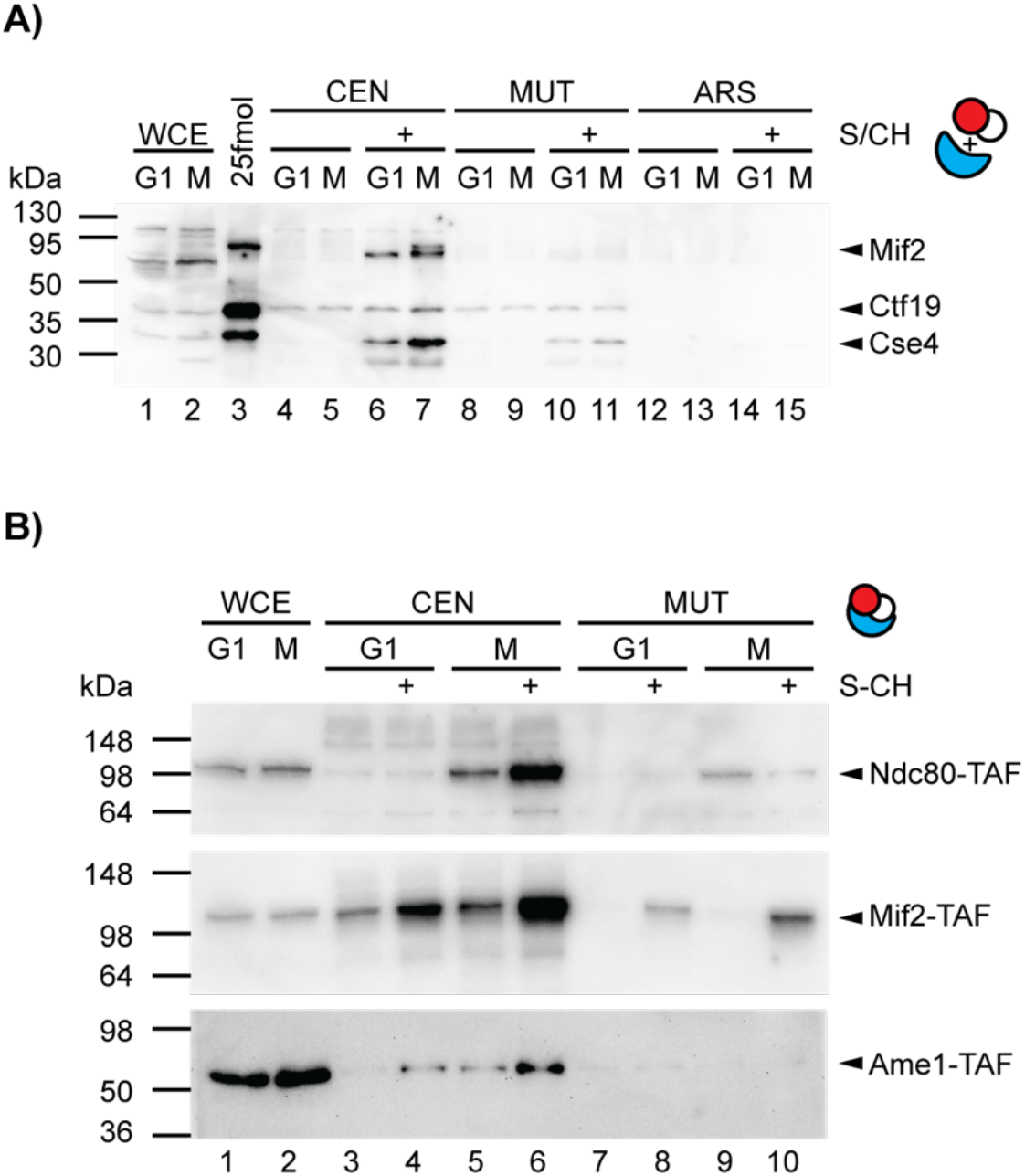
Scm3 promotes cell cycle-dependent kinetochore assembly. (A) Reconstitution in WCEs prepared from cells arrested in G1 with α-factor or in M phase with nocodazole. CEN, MUT, and ARS DNA beads were incubated with the indicated extracts with or without a pre-incubated mixture of separately purified Scm3 and CH (S/CH; 30 nM each). DNA-associated Mif2, Ctf19, and Cse4 were detected by immunoblotting. Lanes 1 and 2 contain G1- and M-phase WCE inputs; lane 3 contains recombinant protein standards; lanes 4-7, 8-11, and 12-15 show CEN, MUT, and ARS reactions, respectively. (B) Reconstitution of inner- and outer-kinetochore components. WCEs were prepared from separate strains expressing 3xFLAG-Protein A (TAF)-tagged Ndc80, Mif2, or Ame1 and arrested in G1 or M phase. CEN and MUT DNA beads were incubated with the corresponding extracts with or without co-purified Scm3/Cse4/H4 (S-CH; 100 nM). DNA-associated tagged proteins were detected using an anti-Protein A antibody. Lanes 1 and 2 contain G1- and M-phase WCE inputs; lanes 3-6 and 7-10 show CEN and MUT reactions, respectively.

To improve detection of additional kinetochore components, reconstitution assays were also performed using G1-and M-phase WCEs prepared from strains expressing Protein A-tagged Ame1, Mif2, or Ndc80, in three separate experiments (**Figure 2B**). SCH addition increased the recovery of Ndc80 and Mif2 on CEN DNA, whereas the increase in Ame1 recovery was more modest, although it remained specific to CEN DNA. For all three proteins, recovery from CEN DNA remained greater in M-phase WCEs than in G1 WCEs. Thus, mitotic cell cycle state enhances the efficiency of Cse4 loading and/or retention on CEN DNA, and all subsequent analysis employed M-phase WCEs.

### The Scm3 interface with Cse4 is essential for stimulation of Cse4 assembly and cell viability

The interaction of Scm3 with the Cse4/H4 dimer has been structurally characterized in the budding yeast *Kluyveromyces lactis*, which diverged from *S. cerevisiae* ∼100 million years ago (51) and shares conserved CDEI and CDEIII elements, although its CDEII element is approximately twice as long as that of *S. cerevisiae* (52). The histone-binding domain of *K. lactis* Scm3 contacts the Cse4/H4 histone dimer through extensive interactions with the long central helix of Cse4’s histone fold domain. Structural and biochemical studies have defined *K. lactis* Scm3 residues responsible for selective Cse4 binding (H67, D71, K84, and Y85 of *K. lactis* Scm3) (31, 49). Sequence alignments showed that these residues are conserved and correspond to H96, D100, K113, and Y114 of *S. cerevisiae* Scm3 (**Figure 3A**); K113 and Y114 are also conserved in human HJURP (49). An AlphaFold model positioned all 4 residues along the α2 helix of Scm3 (**Figure 3B**), consistent with existing structural evidence (31, 49). In a plasmid shuffle assay, mutations of K113 and Y114 to alanines resulted in inviability while mutating H96 and D100 to alanines had no observable effects (**Figure 3C**).

**Figure 3.**
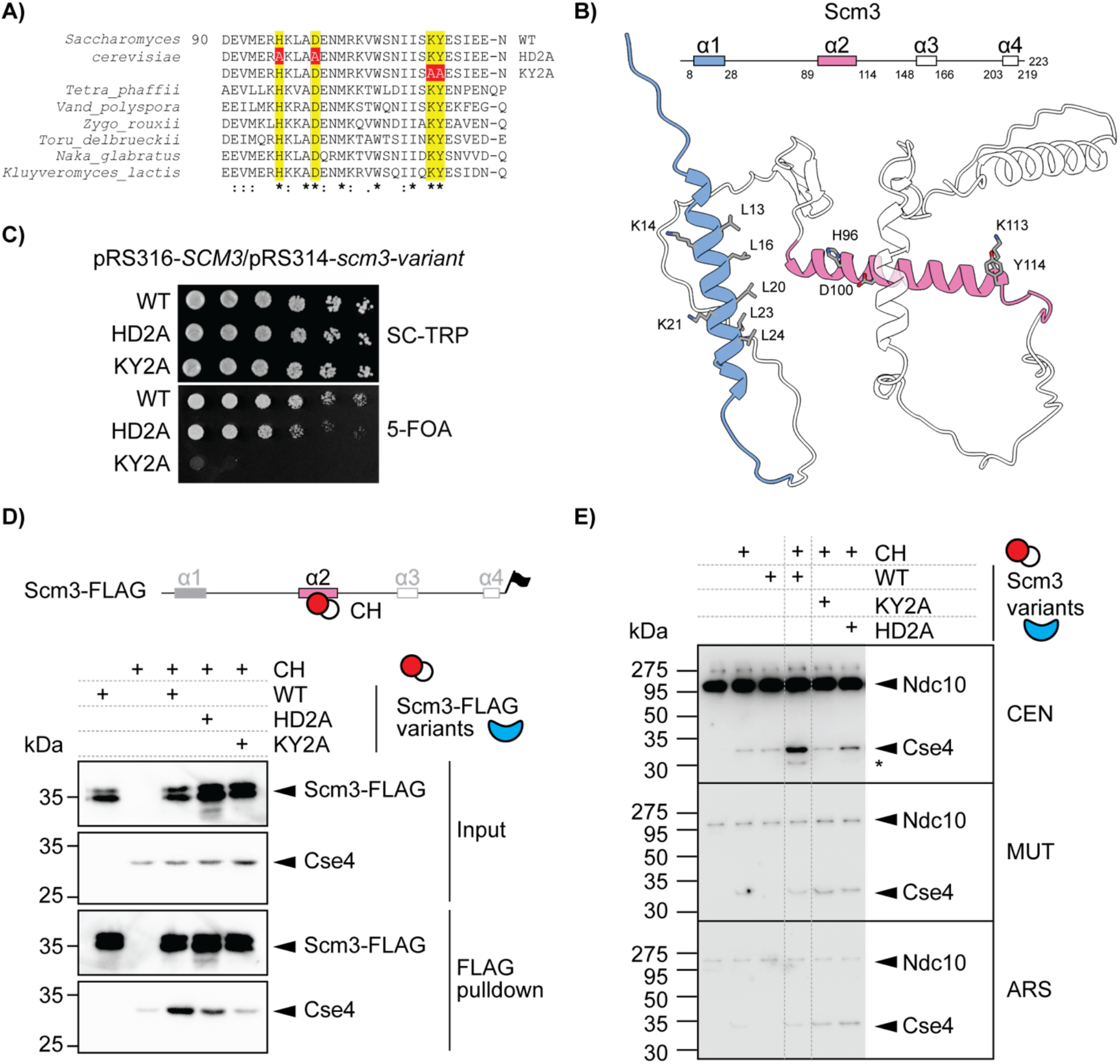
Effects of scm3 mutations at the interface of Scm3 and Cse4/H4. (A) Sequence alignment of the Scm3 histone-binding region. Conserved residues corresponding to H96, D100, K113, and Y114 in *S. cerevisiae* are highlighted in yellow, and alanine substitutions are shown in red. (B) AlphaFold predicted structure and secondary-structure schematic of Scm3. The N-terminal α1 helix and central α2 helix are shown in blue and pink, respectively. Residues selected for alanine substitution are indicated. (C) Plasmid-shuffling analysis of variants. Serial dilutions of *scm3Δ* cells carrying a URA3-marked plasmid expressing WT SCM3 (pRS316-SCM3) and a TRP1-marked plasmid expressing WT or mutant SCM3 (pRS314) were spotted onto synthetic complete medium lacking tryptophan (SC-TRP) or medium containing 5-fluoroorotic acid (5-FOA). Growth on 5-FOA assesses the ability of the indicated variant to support viability following loss of the URA3-marked WT SCM3 plasmid. (D) FLAG pulldown assay of Scm3-Cse4/H4 binding. WT, HD2A, or KY2A Scm3 bearing a C-terminal 6xHis-3xFLAG tag (1 μM) was incubated with Cse4/H4 (1 μM) and captured using anti-FLAG affinity resin. Reactions lacking Scm3 or Cse4/H4 served as controls. Input and bound fractions were analyzed by immunoblotting with anti-FLAG and anti-Cse4 antibodies. Anti-FLAG and anti-Cse4 blots were processed separately. (E) Cell-free reconstitution with WT or mutant Scm3. CEN, MUT, and ARS DNA beads were incubated with WCE alone or supplemented with CH, WT Scm3, or a pre-incubated mixture of CH and WT or mutant Scm3, as indicated. CH and Scm3 were each added at a final concentration of 100 nM where present. DNA-associated Ndc10 and Cse4 were detected by immunoblotting. Asterisks indicate cross-reaction/degradation product.

To determine the significance of the Scm3-Cse4 interface in the Cse4 assembly reaction, we first performed a pulldown assay to examine the binding between recombinant Scm3 (6xHis-3xFLAG, i.e. HF) and CH proteins purified from bacteria. Scm3-HF, along with its mutant variants, were incubated with CH and bound to anti-FLAG affinity beads. Immunoblots against Cse4 revealed that wild-type (WT) Scm3-HF bound to Cse4 (**Figure 3D**) (26, 47, 48), while the HD2A mutant of Scm3 caused mildly lower amounts of co-purified Cse4. The KY2A mutant reduced the co-purified Cse4 to background levels (**Figure 3D**); consistent with observed effects on cell viability (**Figure 3C**).

We next tested the effect of the Scm3 mutations on Cse4 reconstitution in WCE. As predicted, the KY2A mutations abolished the effect of SCH addition on Cse4 assembly (**Figure 3E**). Interestingly, the HD2A mutation also resulted in a significant reduction in Cse4 assembly, likely because mutation of these two residues weakens the Cse4 interface of Scm3 and the stringency of the reconstitution assay reveals this effect. These results establish that the Scm3 interface with Cse4 is essential for the in vitro assembly of Cse4 and for cell viability.

### Interaction of the N-terminus of Scm3 with the CBF3 complex directs Cse4 deposition to native centromeres

The above results indicate that Scm3 recognition of Cse4 is critical for its ability to promote Cse4 assembly at native centromeres. However, they do not address why Scm3 specifically targets Cse4 assembly to WT centromeres. In budding yeast point centromeres, the CDEIII sequence element must bind to the multi-subunit CBF3 complex to initiate kinetochore assembly (25, 38). A direct interaction between the *K. lactis* Scm3 N-terminal domain and the Ndc10 subunit of CBF3, specifically a C-terminal domain of *K. lactis* Ndc10, has been reported (50). This Scm3 N-terminal domain corresponds to the α1 helix of *S. cerevisiae* Scm3 (**Figure 4A** and **3B**). The N-terminal region of Scm3 is required for cell viability (47, 48) and contains a conserved sequence spanning residues 12-24 whose specific function remains undefined (**Figure 3B** and **4A**). To examine whether conserved residues within this sequence contribute to Ndc10 binding, five leucine residues (L13, L16, L20, L23, and L24) and two lysine residues (K14 and K21) were targeted for alanine substitution. Based on an AlphaFold structure prediction, L13, L16, L20, L23 and L24 line one hydrophobic face of the α1 helix, while K14 and K21 sit on the other (**Figure 3B**). Therefore, 5LA removes the whole face, 2LA and 3LA split it, and 2KA targets the basic residues. Employing a plasmid-shuffling assay to replace Scm3 with the variants, we found that the 5LA and 3LA mutants failed to support viability, the 2LA mutant impaired viability, and the 2KA mutant did not significantly affect viability (**Figure 4A** and **4B**). Next, we performed pulldown assays with recombinant WT or mutant variants of the N-terminal domain of Scm3, Scm3^1-60^, and the C-terminal domain of Ndc10, Ndc10^565–956^. The pulldown assays indicated that Ndc10^565-956^ was co-isolated with WT Scm3^1-60^ (**Figure 4C**). The 5LA, 3LA, and 2LA mutations largely disrupted the observed binding between Scm3^1-60^ and Ndc10^565-956^, while the 2KA mutation did not (**Figure 4C**). These in vitro binding assays confirmed a direct Scm3-Ndc10 interaction and defined mutants that perturb the interaction. We next examined the effects of these mutations on Cse4 assembly in the cell-free reconstitution assay (**Figure 4D** and **Supplementary Figure 4**). Addition of WT Scm3 together with CH markedly increased Cse4 assembly on CEN DNA. In contrast, all mutations except for the 2KA abolished the effect of SCH addition, and the 2KA mutant showed a partial yet observable assembly defect. These data indicate that mutations perturbing Scm3’s interface with Ndc10 significantly impair Cse4 assembly on CEN DNA in WCEs.

**Figure 4.**
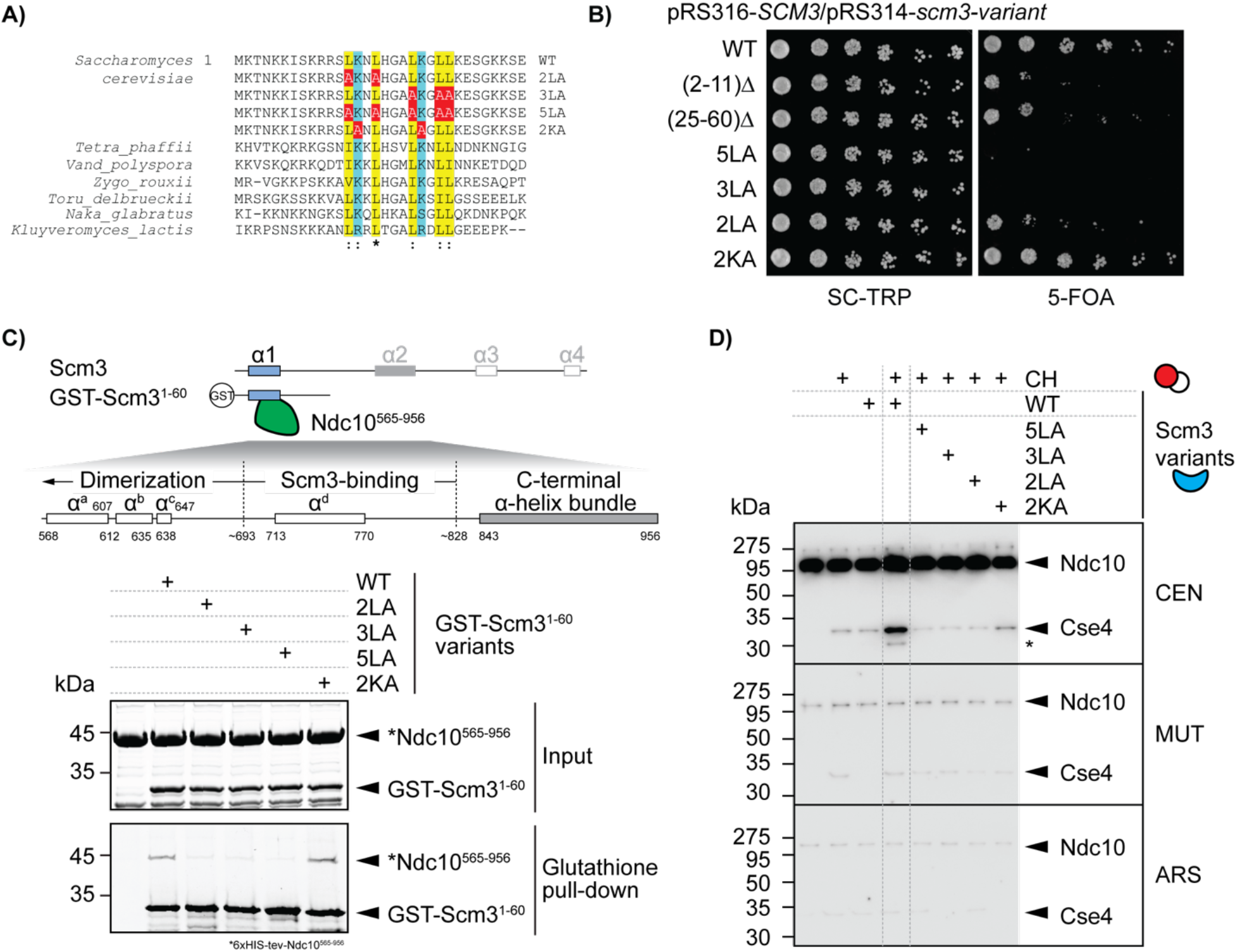
Effects of scm3 mutations on the putative binding interface of Scm3 and Ndc10. (A) Sequence alignment of the putative Ndc10 binding region of Scm3 from *Saccharomyces cerevisiae* and the indicated fungal species, together with the *S. cerevisiae* mutant sequences. Leucine and lysine residues targeted for substitution are highlighted in yellow and cyan, respectively; alanine substitutions are shown in red. Mutants are 5LA (L13A/L16A/L20A/L23A/L24A), 3LA (L20A/L23A/L24A), 2LA (L13A/L16A), and 2KA (K14A/K21A). (B) Plasmid-shuffling analysis of 2 N-terminus truncation mutants of Scm3 (Δ2-11 and Δ25-60) and variants carrying point mutations at the putative Ndc10 binding domain (5LA, 3LA, 2LA, and 2KA), performed as described in panel 3C. (C) Top: Domain schematics of the Ndc10 C-terminal fragment (residues 565-956) and Scm3, indicating the regions potentially involved in Scm3-Ndc10 binding. Bottom: GST pulldown assay. Recombinant 6xHis-tagged Ndc10^565-956^ (10 μM) was incubated with GST-Scm3^1-60^ (2 μM), containing the WT sequence or the indicated 2LA, 3LA, 5LA, or 2KA substitutions, followed by capture on glutathione agarose resin. Input and bound fractions were analyzed by SDS-PAGE followed by Coomassie Brilliant Blue staining. The control reaction contained Ndc10 without GST-Scm3. Protein positions are indicated; the asterisk specifies that Ndc10^565-956^ has an N-terminal 6xHIS tag. (D) Cell-free reconstitution with WT or mutant Scm3, performed as described in panel 3E.

Considering that Ndc10, as a part of the CBF3 complex, resides at centromeres in cells, these results suggest that the N-terminal mutants of Scm3 that disrupt Ndc10 binding should also impair Scm3’s localization to CEN DNA in vivo. To test this hypothesis, we expressed Flag-tagged WT and N-terminal mutants of Scm3 from a centromeric plasmid under its native Scm3 promoter and then conducted ChIP-qPCR analysis. We found that 5LA, 3LA and 2LA mutants have significantly reduced CEN association (**Figure 5A**), while the 2KA mutant shows a slightly reduced CEN-association compared to WT Scm3. This finding agrees with the effects of these mutants on the binding between Scm3 N-terminus and Ndc10 (**Figure 4C**), indicating that the Scm3-Ndc10 interaction is responsible for recruiting Scm3 to centromeres in vivo.

**Figure 5.**
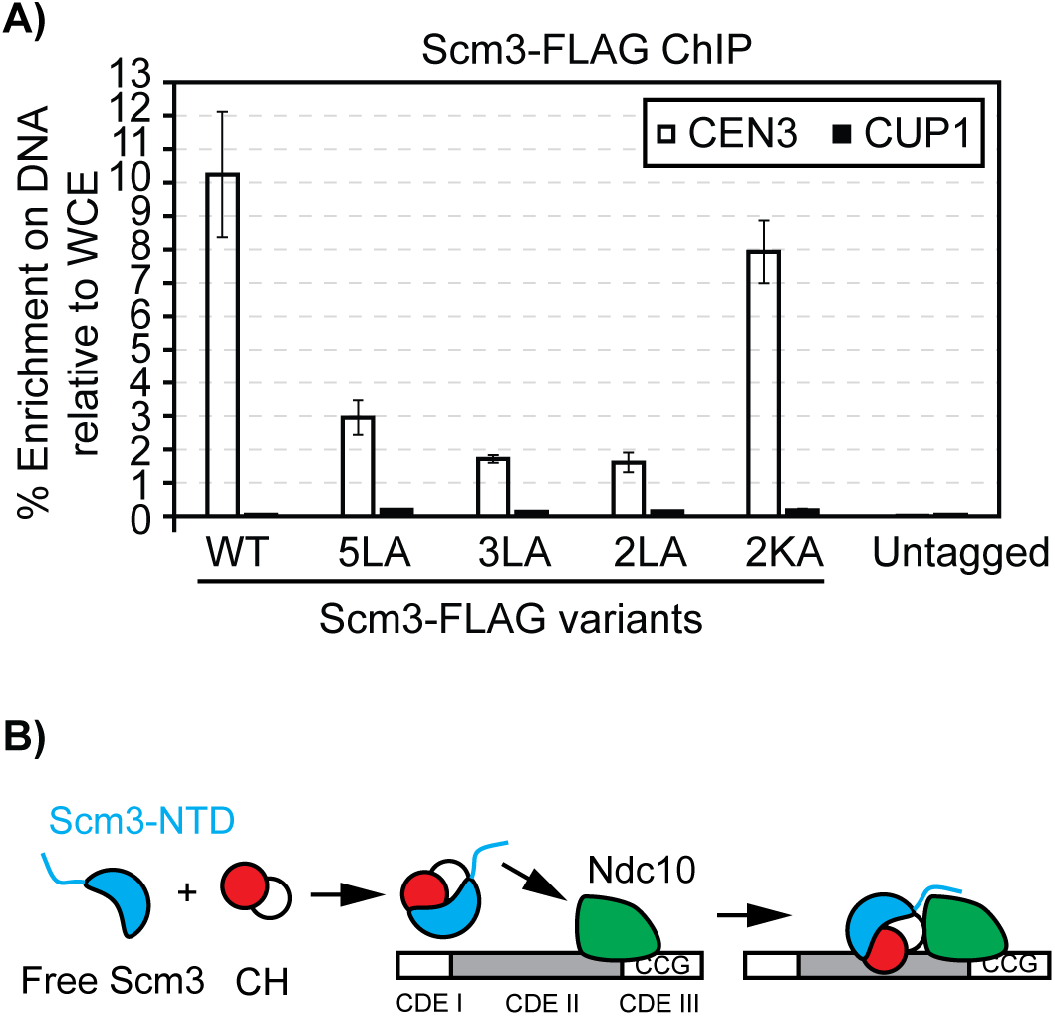
Scm3 interface with Ndc10 is important for Scm3 centromere localization in vivo. A) Chromatin immunoprecipitation followed by quantitative PCR (ChIP-qPCR) of plasmid-expressed, 3xFLAG-tagged WT and mutant Scm3. Open bars show enrichment at centromere III (CEN3), and filled bars show enrichment at the CUP1 promoter control region. Enrichment is expressed as the percentage of input DNA measured in WCE. An untagged strain served as a negative control. Bars show means, and error bars indicate standard deviations from three parallel immunoprecipitations performed on the same cell lysate for each strain. B) Schematic model of Scm3-dependent Cse4 assembly on CEN DNA. Scm3 binds Cse4/H4 and engages centromere-associated Ndc10 through its N-terminal domain (Scm3-NTD), promoting recruitment and assembly of Cse4/H4 on centromeric DNA.

Taken together, these results show that an essential function of the conserved Scm3 N-terminus is to contact Ndc10, allowing the recruitment of Scm3 to the centromeres where Scm3 promotes Cse4 assembly.

## DISCUSSION

Assembly of Cse4^CENP-A^ containing kinetochores on native centromeric DNA is a prerequisite for building the kinetochore. Many cellular factors have been reported to regulate the assembly, but their mechanisms of action remain poorly understood. Using a cell-free system from yeast extracts, we show that Scm3 is a limiting factor for Cse4 assembly on native centromeric DNA. Adding Scm3 together with Cse4/H4 strongly stimulated Cse4 assembly in a DNA sequence- and dose-dependent manner (**Figure 1**). It also increased recruitment of downstream inner and outer kinetochore proteins throughout the cell cycle (**Figure 2**). This stimulation required two distinct Scm3 interfaces: the α2-helix surface that binds Cse4/H4, and the N-terminal α1 helix that binds the C-terminal domain of Ndc10 (**Figure 3** and **Figure 4**). We propose that Scm3 delivers Cse4/H4 to the centromere by engaging Ndc10 within the CDEIII-bound CBF3 complex (**Figure 5B**). Thus, Scm3 links sequence-specific centromere recognition to centromeric histone deposition.

Collectively, the genetic and biochemical findings here support the function of Scm3 in specifically promoting Cse4 assembly on native centromeres. It should also be noted that, although the extent of the *scm3* mutant defects varies across different assays, likely due to the in vitro assembly assay being the most stringent, the overall trend consistently supports this model (**Figure 5B**). For example, the 5LA, 3LA and KY2A mutants are inviable while showing strong biochemical defects. The 2LA and HD2A mutants show minimal growth defects while showing clear defects in vitro. The 2KA mutant retained Ndc10 binding but showed a noticeable defect in Cse4 assembly and a mild reduction of Scm3 localization at the centromere in cells, consistent with its lack of appreciable growth defect (**Figure 4B**).

Our results have multiple implications and point to future directions. First, the dose dependence study suggests that cellular levels of Scm3 and Cse4/H4 must be tightly controlled to prevent non-centromeric Cse4 deposition (**Figure 1D**). This is consistent with the observation that Cse4 overexpression causes its incorporation outside centromeres and impairs growth of cells lacking the ubiquitin ligase Psh1 (53–55). Thus, future attempts to reconstitute Cse4 assembly with purified proteins should aim at meeting this strict centromeric DNA sequence requirement and understanding its peculiar sequence characteristics.

Second, adding SCH promoted kinetochore assembly in both G1- and M-phase extracts but did not eliminate the difference between them: assembly was consistently stronger in M-phase extracts (**Figure 2**). This points to additional cell cycle control of inner kinetochore assembly, consistent with mechanisms of phospho-regulation of inner and outer kinetochore assembly (1, 34, 56–58). In yeast, Cse4 resides at centromeres for most of the cell cycle and turns over briefly during centromere replication (59); in human cells, new CENP-A is deposited in early G1 (60). Regulated destabilization of centromeric histone assembly may therefore be needed to accommodate centromere replication. Viewed from this lens, the weaker Cse4 assembly in G1 extracts may reflect a more dynamic equilibrium, while additional M-phase specific signal(s) would strengthen the assembly in preparation for chromosome segregation. Although the recombinant SCH produced in bacteria lacks post-translational modifications, it may be modified in M-phase extracts, or it may interact with partners that are modified in M-phase. Alternatively, destabilizing factors may act specifically in G1, while stabilizing factors may act in the M-phase. Further studies are needed to distinguish these possibilities.

Finally, our analysis of the assembled products from the cell-free reconstitution is an endpoint analysis. It therefore cannot distinguish assembly (the forward reaction, such as Scm3 stimulation) from stabilization by other proteins or from passive or active disassembly, all of which may proceed simultaneously in the extracts. For instance, a less stable Cse4-CEN association in G1 extracts would be indistinguishable from reduced loading. In addition, the Scm3–Ndc10 interaction was assayed with an N-terminal fragment of Scm3 and a C-terminal fragment of Ndc10, rather than with full-length Scm3 and the intact CBF3 complex and may not fully reflect the native interaction. Nevertheless, establishing Scm3 as a key driver of Cse4 assembly on native centromeric DNA demonstrates the utility of this approach for dissecting the factors that contribute to kinetochore assembly and for understanding how this fundamental process operates.

## MATERIALS and METHODS

### Yeast genetics method and plasmid construction

Standard yeast genetics methods were used to construct the strains used in this study. Unless noted otherwise, plasmids were constructed using DNA recombination repair in yeast via >35 base pairs of homologies among PCR fragments used. Plasmids were rescued from yeast via electroporation, purified from bacteria and confirmed by DNA sequencing. Details of these constructions are available upon request. All plasmids and yeast strains used in this study can be found in **Supplementary Table 1A** and **1B**. Plasmid shuffling was performed as previously described (61).

### Spotting assay

Yeast cells were grown in 5 mL of SC-TRP media at 30 °C until saturation. Cultures were normalized to an OD_600_ of 1.0, followed by 5-fold serial dilutions using multichannel pipettes in a 96-well plate. From each dilution, 8 μL of cells were spotted on SC-TRP or 5-FOA media. Plates were incubated at 30 °C for 3 days and then imaged using a Bio-Rad ChemiDoc imaging system.

### Yeast cell extract preparation for cell-free kinetochore reconstitution

Yeast cells (HZY1029 and others, see **Supplementary Table 1**) were grown in 1 L of YPD (yeast extract, peptone, and dextrose) medium at 30 °C to OD_600_ of 0.3 and arrested in G1- or M-phase by adding 15 nM alpha factor or 15 μg/mL nocodazole, respectively, for 3 hours. Completion of the arrests was confirmed by cell morphology. Cells were centrifuged at 4,000 RCF for 8 minutes, and the resulting cell pellet was washed once with PBS (pH 8.0) and resuspended in 1/4 cell pellet volume buffer L (25 mM HEPES-KOH, pH 8.0/175 mM K-glutamate/2 mM MgCl_2_/0.1 mM EDTA/0.5 mM EGTA/0.1% NP-40/15% glycerol/protease inhibitor cocktail (PI) (Sigma-Aldrich, Cat #: P-8849)/1 mM Phenylmethylsulfonyl fluoride (PMSF) in deionized H_2_O). This yeast cell resuspension was flash-frozen in liquid nitrogen drop by drop (popcorn) and stored at -80 °C. Before the reconstitution experiments, frozen cell popcorn was lysed using a cryogenic grinder (Cat #: 6875D115; SPEX SamplePrep LLC) at 10 cycles per second for 12 cycles, with a 2-min cycle time and a 2-min cooling time between cycles. The lysed cell powder was thawed on ice and centrifuged at 21,000 RCF for 30 minutes, and the clear fraction was separated from the pellet, aliquoted, and stored at -80 °C until use. The protein concentration of the clarified yeast extract was ∼50 mg/mL, as determined by Bradford assay (Bio-Rad).

### Preparation of centromeric and control DNA beads

Yeast centromere III (CEN), centromere III with point mutations to its CDE III region (MUT), and yeast autonomously replicating sequence (ARS) were amplified from the plasmids HZE3240, HZE3241, and HZE3246, respectively, under standard PCR conditions with 0.5 μg Taq polymerase (in-house-purified)/20 mM Tris-HCl (pH 8.4)/50 mM KCl/dNTP (50 μM each)/2 mM MgCl_2_ in deionized H_2_O and primers 1 and 2 for CEN and MUT DNA, or primers 3 and 4 for ARS DNA (**Supplementary Table 1**).

To improve PCR efficiency, extra dTTP and dATP were added (to a final concentration of 200 μM each) when amplifying CEN and MUT DNA. The resulting PCR product was 230 bp in length. PCR products were precipitated and washed with 100 mM ammonium acetate in 70% ethanol/30% deionized H_2_O. Following precipitation, the DNA pellet was resuspended in 10 mM Tris-HCl/1 mM EDTA (pH 7.5) and purified using the Wizard SV PCR Clean-up Kit (Cat #: A9281, Promega). The concentrations of the purified DNA were measured by a NanoDrop spectrophotometer (Cat #: ND2000; Thermo Fisher Scientific). Before each reconstitution, 200 μL of Dynabeads M-280 streptavidin (Cat #: 11206D; Thermo Fisher Scientific) was incubated with 10 μg of the purified DNA at room temperature for 20minutes in 1 M NaCl/0.5 mM EDTA/5 mM Tris-HCl (pH 7.5) in deionized H_2_O. The amount of DNA bound to the beads was calculated to be ∼4 μg per 100 μL of the original bead solution. After binding with DNA, the beads were washed twice with buffer L and stored at -20 °C until use.

### Purification of recombinant SCH, Cbf1, Sgt1, Scm3^1-60^, and Ndc10^565-956^

All recombinant proteins were expressed in *E. coli* Rosetta (DE3) cells, transformed by various plasmids (**Supplementary Table 1)**. *E. coli* cells were grown at 37 °C to OD_600_ of 0.6 before induction with 0.1 mM isopropyl β-D-1-thiogalactopyranoside (IPTG) at 17 °C overnight. Cells were harvested by centrifugation (10 minutes at 4,000 RCF) and resuspended in PBS buffer supplemented with PI and 1 mM PMSF. Cell pellets were stored at -80 °C until purification. Cell lysis was performed by sonication, and cell extracts were clarified by centrifugation at 21,000 RCF for 30 minutes. 6xHis-tagged proteins were purified by Ni-NTA affinity resins (Qiagen, Cat #: 30210), washed with PBS buffer supplemented with 0.5 M NaCl, 0.1% NP-40, and 20 mM imidazole, and eluted in PBS buffer supplemented with 0.5 M NaCl, 0.1% NP-40, and 200 mM imidazole. The eluates were adjusted to a final glycerol concentration of 5% (v/v) and stored at - 80 °C until use.

For the purification of SCH constructs and Cse4/H4 constructs as complexes, Ni eluates were loaded on a Superdex 200 Increase 10/300 GL SEC column (Cytiva 28-9909-44, column L x I.D. 30 cm x 10 mm, 8.6 μm particle size) and purified at a flow rate of 0.5 mL/min with a high-salt (HS) buffer (0.5 M NaCl/5 mM β-mercaptoethanol/PBS). Eluates from the column were collected into 1 mL fractions and evaluated by UV absorption at 280 nm and Coomassie Brilliant Blue-stained Sodium Dodecyl Sulfate-Polyacrylamide Gel Electrophoresis (SDS-PAGE) to visualize the desired proteins. The fractions with the highest protein content were pooled, concentrated using centrifugal filter units (Amicon Ultra - 0.5 mL Ultracel 10K NMWL, Millipore, Cat #: UFC501024), and glycerol was added to the pooled eluates to a final concentration of 5% (v/v). Aliquots of the concentrated eluates were then stored at -80 °C until use. All recombinant SCH and Cse4/H4 constructs, along with BSA standards, were visualized by Coomassie Brilliant Blue-stained SDS-PAGE. (**Supplementary Figure 1** and **Supplementary Figure 4**).

GST-tagged Scm3^1-60^ variants were purified using Pierce Glutathione Agarose resin (Thermo Scientific, Cat #: 16101) following standard protocol. Briefly, Rosetta cells carrying plasmids expressing GST-tagged Scm3^1-60^ variants were induced and lysed. Centrifugation clarified cell lysates were transferred to glutathione resin and incubated at 4 °C for 60 minutes. The resin was subsequently washed with PBS buffer supplemented with PI, and bound proteins were eluted by TNGT buffer (50 mM Tris-HCl, 100 mM NaCl, 0.2% NP-40, 50 mM reduced glutathione, pH 8.33). Glycerol was added to the eluate to a final concentration of 10% (v/v), and the eluates were stored at -80 °C until use.

### Cell-free reconstitution of kinetochores

Frozen yeast extracts were thawed from -80 °C on ice before reconstitution experiments. For each kinetochore reconstitution, 10 μL of DNA beads were incubated with 100 μL of yeast extract at room temperature for 1.5 hours. After incubation, the beads were washed four times with 500 μL ice-cold buffer L supplemented with protease inhibitors. To elute reconstituted kinetochores, the washed beads were resuspended in 30 μL DNase I buffer (50mM HEPES-KOH pH 7.45/10mM MgCl_2_/10mM CaCl_2_/0.1 M NaCl/0.1% NP-40/1mM EDTA pH 7.45/PI) with 0.1 U/μL TURBO DNase I (Cat #: AM2238, Thermo Fisher Scientific) and incubated at room temperature for 1 hr. The DNase-I eluted samples were collected for analysis. Alternatively, the washed beads were directly stripped by 1% SDS at 65 °C for 10 minutes, no differences were observed between the two elution strategies.

For reconstitution experiments involving the addition of recombinant proteins, recombinant proteins were pre-incubated in small volumes on ice for 20 minutes before being added to the yeast lysates. SCH supplements are either purified as a complex or a product of a pre-incubated mix of separately purified Cse4/H4 and Scm3. Final concentrations and method of addition (pre-incubation or as complex) of each recombinant protein/protein complex are shown in **Supplementary Table 2**.

### Pulldown assays to examine interactions between Scm3, Cse4/H4 and Ndc10

To detect binding between Scm3 and Ndc10, 2 μM of GST-tagged Scm3^1-60^ variants and 10 μM of 6xHis-tagged Ndc10^565-956^ were preincubated in 100 μL of binding buffer (1xPBS/0.2% NP-40/10% (v/v) glycerol) for 10 minutes at 4 °C with rotation. The protein mixture was then incubated with 20 μL glutathione agarose resin (GoldBio, catalog no. G-250-10) for 2 hours at 4 °C with rotation. The flow-through was discarded, and the resin was washed seven times with 1 mL of the binding buffer. Bound proteins were eluted with 50 μL of 1% SDS/10 mM dithiothreitol (DTT).

To detect binding between Scm3 and Cse4, 1 μM of 3xFLAG-tagged Scm3 variants and 1 μM of Cse4/H4 were preincubated in 100 μL of binding buffer for 10 minutes at 4 °C with rotation. The protein mixture was then incubated with 20 μL Anti-FLAG M2 affinity resin for 2.5 hours at 4 °C with rotation. The flow-through was discarded, and the resin was washed seven times with 1 mL of binding buffer. Bound proteins were eluted with 50 μL of 1% SDS/10 mM DTT.

### Immuno-blotting

Samples were prepared as described above and separated by SDS-PAGE. Proteins were transferred to a PVDF membrane (BioRad), and immunoblotting was performed using standard protocols (62). Primary antibodies used are shown in **Supplementary Table 3**. Rabbit polyclonal antibodies raised against Ndc10, Cse4, Mif2, Ctf19, and Ndc80 have been described in other studies (34). Goat Anti-Rabbit IgG HRP Conjugate (Millipore, Cat #: 120348) was used as a secondary antibody and visualized by SuperSignal West Dura Extended Duration Substrate and ECL Western Blotting Ultimate Sensitivity Substrate (Femto) (YamayBio, Cat #: VQ4726). Chemiluminescence was recorded on the ChemiDoc Imaging System (Bio-Rad, Cat #: 12003153). Immunoblot exposures vary to best represent differences across samples. All results presented within a subfigure are generated from multiple gels that were transferred simultaneously onto a single PVDF membrane, which was subsequently blocked, incubated with antibodies, washed, and exposed for consistent results across the same blot; therefore, within each subfigure, panels can be compared with each other, with the exception of **Figure 1C** and **3D**. In **Figure 1C**, different exposure times were picked for Ndc10 and Cse4. In **Figure 3D**, anti-FLAG immunoblot was done separately from the anti-Cse4 immunoblot.

Densitometric analysis was performed using custom-developed software to quantify Ndc10 and Cse4 band intensities in the immunoblots shown in **Figure 1D**. Regions of interest were defined around each band, and background subtraction was applied to obtain corrected signals. Negative values resulting from background subtraction were retained. Background-corrected signals were expressed in arbitrary units and plotted against the concentration of added Scm3/Cse4/H4 for each DNA substrate (**Figure 1E**). Quantification was restricted to the immunoblots shown in **Figure 1D**.

### ChIP-qPCR analysis of Scm3

To evaluate the localization of Scm3 and Scm3 variants to the centromere, ChIP-qPCR was performed as previously described (63). Briefly, plasmids expressing Scm3 variants with C-terminal 3xFLAG tags (HZE2450, 2451, 2452, 2453, 2454, 2480, and 2481) were transformed into HZY906 (**Supplementary Table 1**, HZY906 contains another plasmid expressing the WT Scm3 to maintain cell viability); yeast cultures were grown to an OD_600_ of 0.8 in SC-TRP-URA double drop-out media and cross-linked for 15 minutes with 1 % formaldehyde at room temperature. The crosslinked cells were subsequently quenched with glycine, washed, and harvested. Whole-cell lysates were prepared in 0.8 mL ChIP lysis buffer (50 mM HEPES, pH 7.6/140 mM NaCl/1 mM EDTA/1 % Triton/0.1% sodium deoxycholate/PI) by glass bead beating and sonication. Immunoprecipitation (IP) was performed in triplicates (3 process replicates from the same cell lysates) using 50 μL Dynabeads (Protein G; ThermoFisher) and 3 μL anti-FLAG antibody M2 (Sigma-Aldrich; F3165) per IP. After binding, beads were washed, and bound material was eluted, treated by RNase A and proteinase K, and reverse crosslinked at 65 °C overnight. After the reverse crosslinking, DNA was purified using a QIAquick PCR Purification kit (QIAGEN). qPCR was done using PerfeCTa SYBR Green FastMix (QuantaBio, Cat #:95074-012) on a BIO-RAD CFX Opus 96 system. Primers used for qPCR are listed in **Supplementary Table 1** (primers 5, 6, 7, and 8).

### AlphaFold structure and multiple sequence alignment (MSA)

AlphaFold model of Scm3 was obtained from AlphaFold Protein Structure Database (Uniprot: Q12334). The relevant amino acids and secondary structures were visualized using ChimerX (64). MSA was performed using EMBL-EBI Clustal Omega (65).

## Author Contributions

Jiaxi Cai: Conceptualization, data curation, formal analysis, investigation, methodology, resources, software, supervision, validation, visualization, writing - original draft, writing - review & editing.

Ziyi Wang: Data curation, investigation, methodology, resources, visualization, writing - review & editing.

Lily Cheng: Data curation, investigation, methodology, resources. Tianjian Xu: Data curation, investigation.

Cindy Zhang: Data curation, visualization, investigation.

Stephen M. Hinshaw: Conceptualization, investigation, methodology, project administration, resources, supervision, visualization, writing - review & editing.

Raymond T. Suhandynata: Conceptualization, funding acquisition, investigation, resources, supervision, visualization, writing - review & editing

Arshad Desai: Conceptualization, funding acquisition, investigation, methodology, project administration, resources, supervision, visualization, writing - review & editing

Huilin Zhou: Conceptualization, data curation, formal analysis, funding acquisition, investigation, methodology, project administration, resources, software, supervision, validation, visualization, writing - original draft, writing - review & editing.

## DATA AND CODE AVAILABILITY

The densitometry software is available at [GitHub/Zenodo: https://github.com/JiaxiCai/western-blot-densitometry; DOI: 10.5281/zenodo.23253141]. Additional data are available from the corresponding author upon request.

## ACKNOWLEDGEMENTS

This work was supported by the National Institutes of Health through the following grants: GM151191 (Suhandynata, Desai, and Zhou) and GM156288 (Zhou). Jiaxi Cai was supported by a graduate student fellowship from Pfizer-Cell Signaling San Diego.

## SUPPLEMENTARY MATERIAL AND METHODS

**Supplementary Figure 1.**
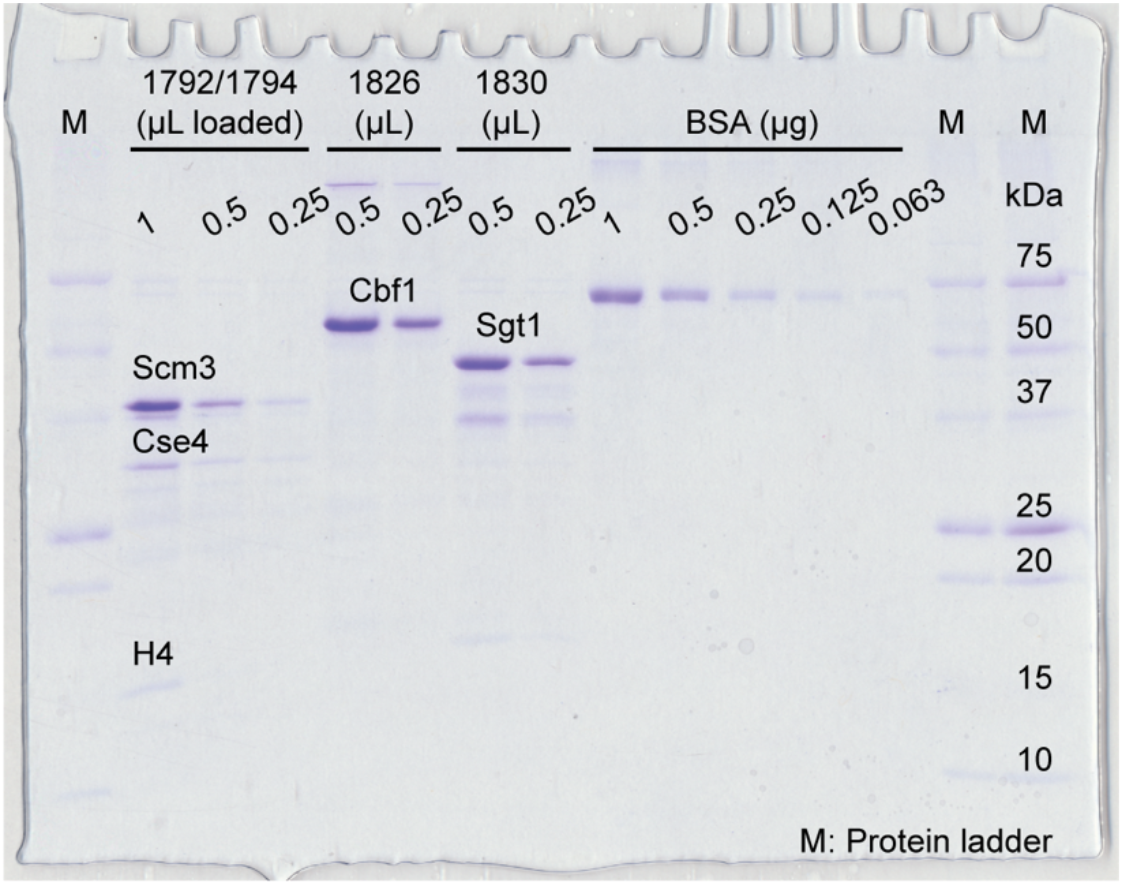
Purified recombinant Scm3/Cse4/H4, Sgt1, and Cbf1. Co-purified Scm3/Cse4/H4, Cbf1, and Sgt1 preparations were analyzed by Coomassie Brilliant Blue-stained SDS-PAGE alongside serial dilutions of bovine serum albumin (BSA) standards. Plasmid identifiers and volumes of recombinant protein preparations loaded are indicated above the lanes; BSA amounts are given in micrograms. Protein identities and molecular masses are indicated. M, molecular mass marker. Protein concentrations used in the reconstitution assays were estimated by comparing band intensities with the BSA standards and are listed in **Supplementary Table 2**.

**Supplementary Figure 2.**
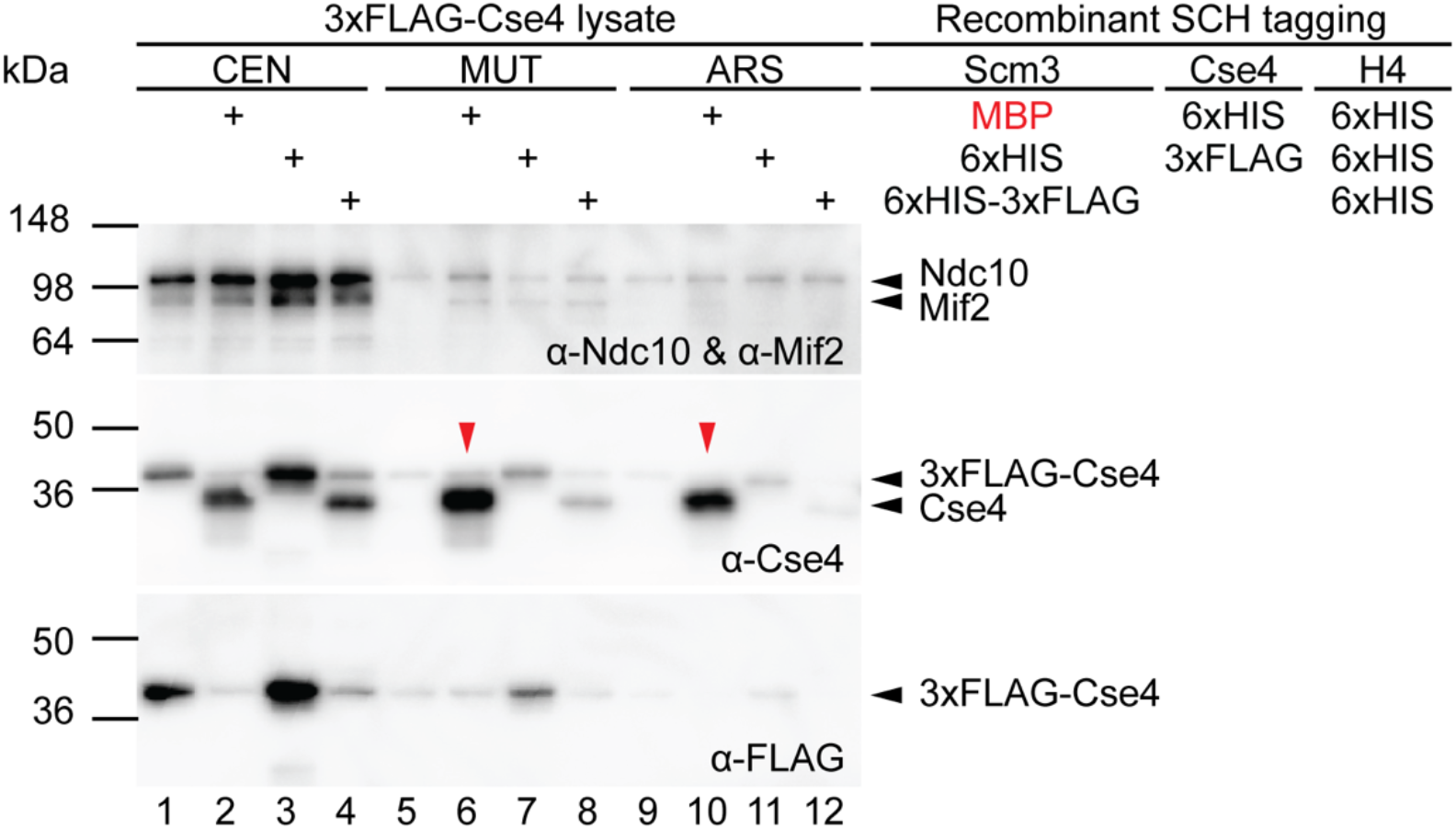
Epitope tagging of SCH can affect its ability to assemble kinetochores. CEN, MUT, and ARS DNA beads were incubated with WCE from strain HZY2777, which expresses endogenous Cse4 containing an internal 3xFLAG tag, with or without recombinant Scm3/Cse4/H4 complexes bearing the indicated epitope tags (100 nM). DNA-associated Ndc10 and Cse4 were detected by immunoblotting. The recombinant preparations included MBP-tagged Scm3 with 6xHis-tagged Cse4 and H4, 6xHis-tagged Scm3 and H4 with internally 3xFLAG-tagged Cse4, and C-terminally 6xHis-3xFLAG-tagged Scm3 with untagged Cse4 and 6xHis-tagged H4. The migration difference between endogenous 3xFLAG-tagged Cse4 and untagged recombinant Cse4 permits their distinction in the corresponding reactions. MBP, maltose-binding protein. The SCH variant containing MBP (maltose-binding protein)-Scm3 and 6xHIS-Cse4 caused elevated Cse4 on all three DNA substrates. By contrast, the two other SCH variants led to CEN-specific increases in Cse4 recovery. Moreover, the addition of the SCH variant with untagged Cse4, which migrates faster than the endogenous 3xFLAG-Cse4 in the gel, caused a specific increase in its binding to CEN-DNA with a simultaneous reduction in the binding of endogenous 3xFLAG-Cse4, suggesting that both 3xFLAG-tagged and untagged versions of Cse4 can be assembled in a CEN-specific manner. In all cases, the addition of these SCH variants did not affect the reconstitution of Ndc10. Red arrows indicate non-CEN association of Cse4 when an MBP tag on Scm3 was present.

**Supplementary Figure 3.**
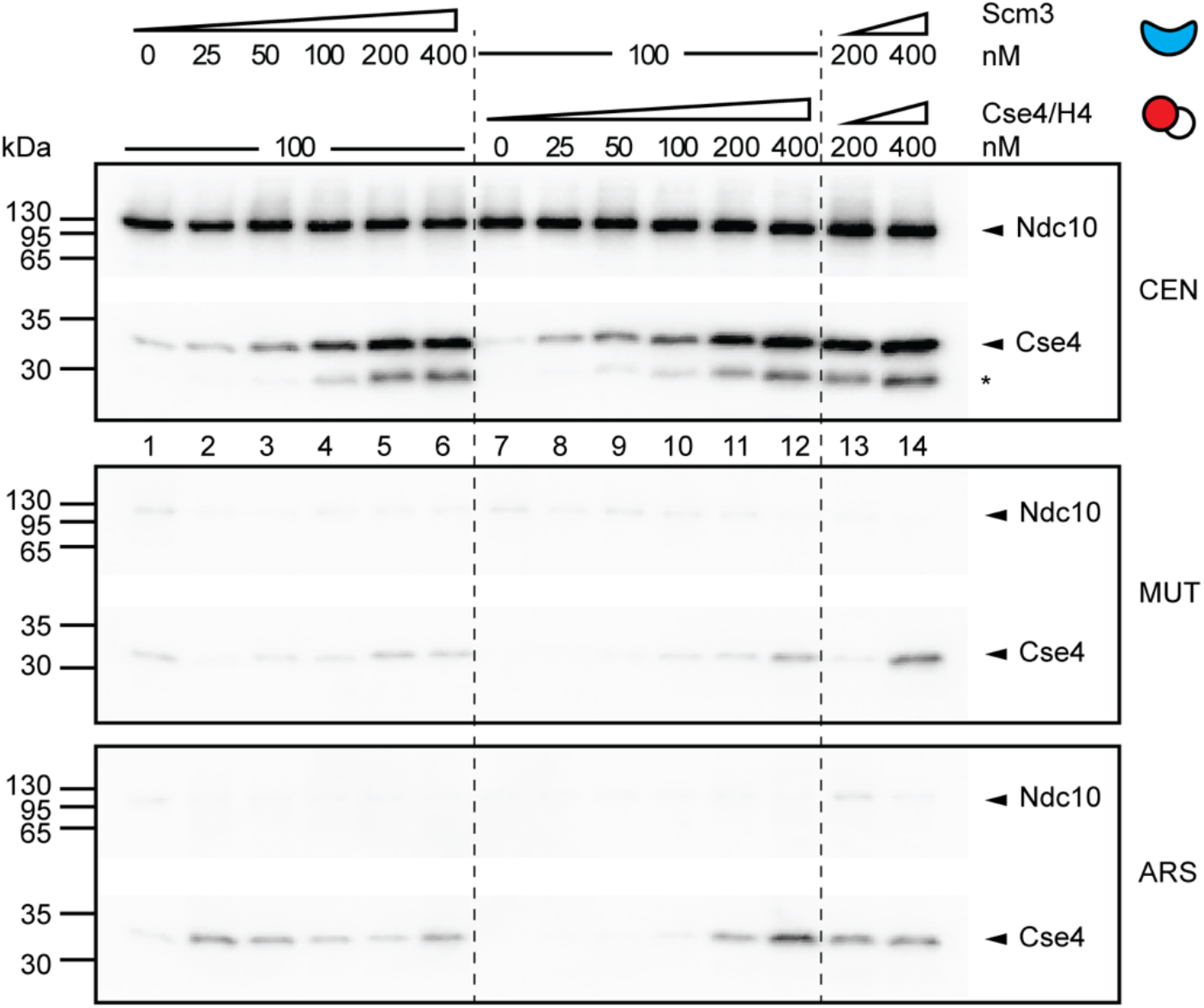
Titration of Cse4/H4 and Scm3 both led to improved Cse4 reconstitution. CEN, MUT, and ARS DNA beads were incubated with WCE supplemented with pre-incubated mixtures of separately purified Scm3 and Cse4/H4 (CH). In lanes 1-6, Scm3 was titrated from 0 to 400 nM with CH held at 100 nM. In lanes 7-12, CH was titrated from 0 to 400 nM with Scm3 held at 100 nM. Lanes 13 and 14 contain both Scm3 and CH at 200 and 400 nM each, respectively. DNA-associated Ndc10 and Cse4 were detected by immunoblotting. Dashed vertical lines separate the titration series.

**Supplementary Figure 4.**
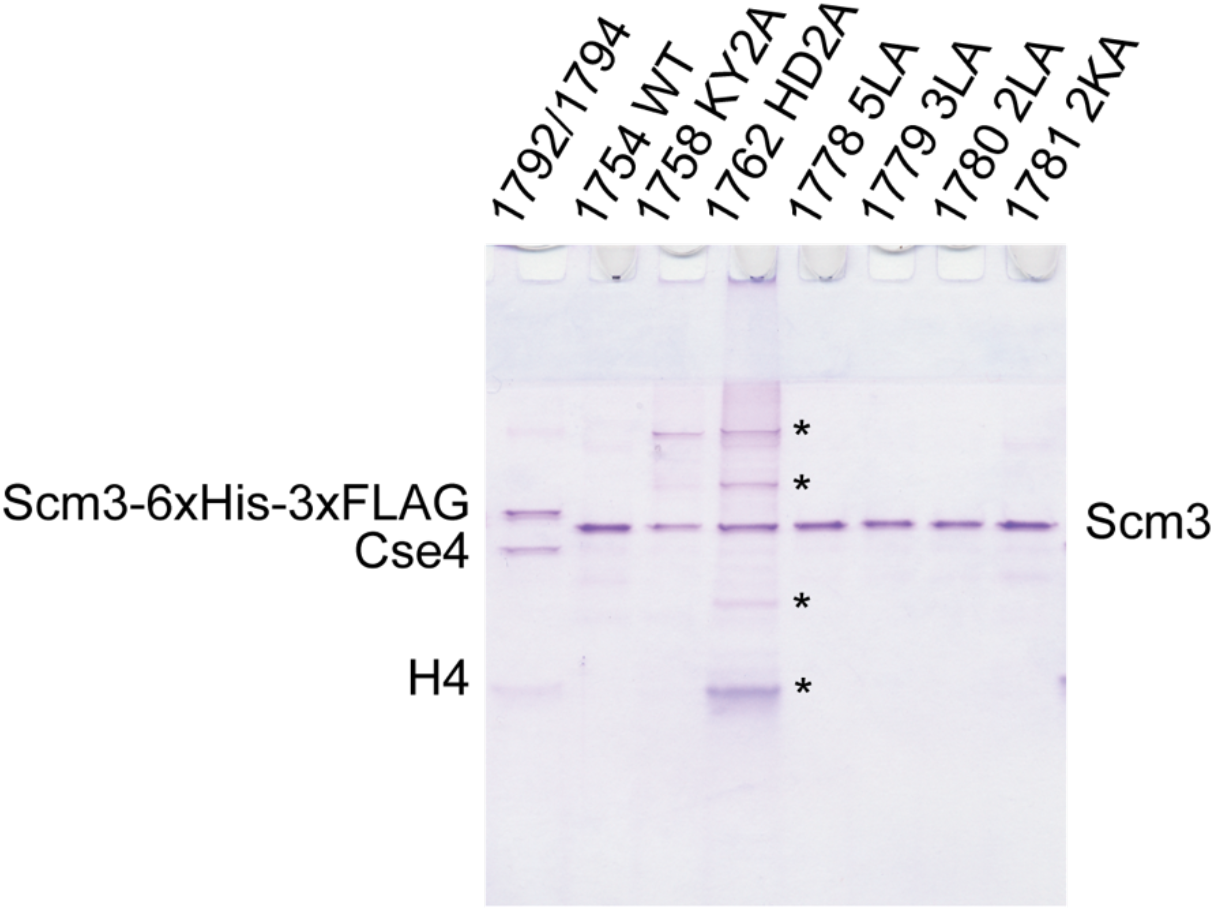
Purified Scm3 proteins used in Figure 4 to study the effect of scm3 mutations for Cse4 loading. Recombinant WT and mutant Scm3 preparations were analyzed by Coomassie Brilliant Blue-stained SDS-PAGE. Plasmid identifiers and Scm3 variants are indicated above the lanes. The first lane contains Cse4/H4 and WT Scm3 preparations (plasmids HZE1792 and HZE1794 were co-expressed and proteins were co-purified); the remaining lanes contain WT Scm3 or the KY2A, HD2A, 5LA, 3LA, 2LA, and 2KA variants. Protein positions are indicated. Asterisks mark additional protein species in the HD2A preparation.

**Supplementary Table 1.**
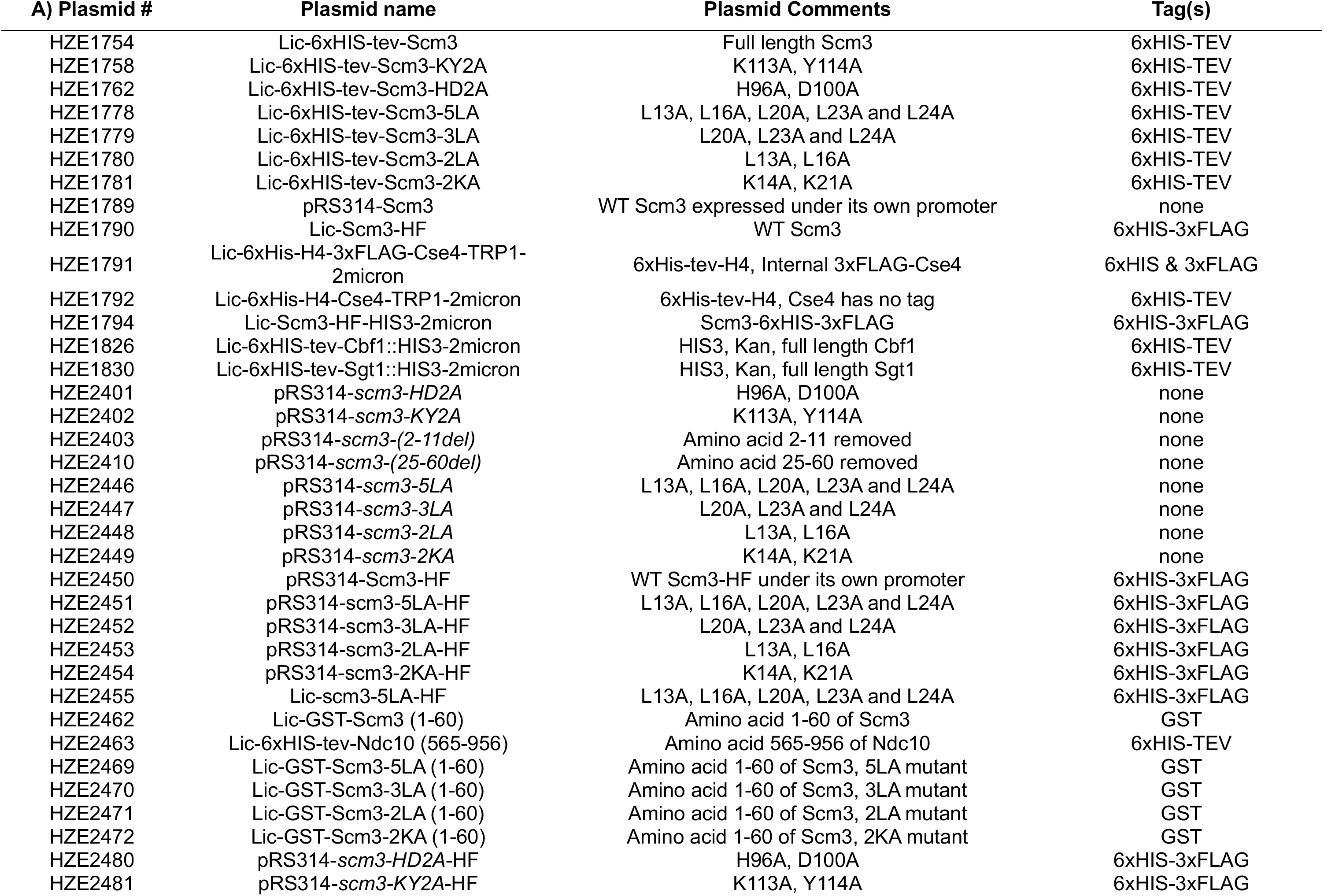
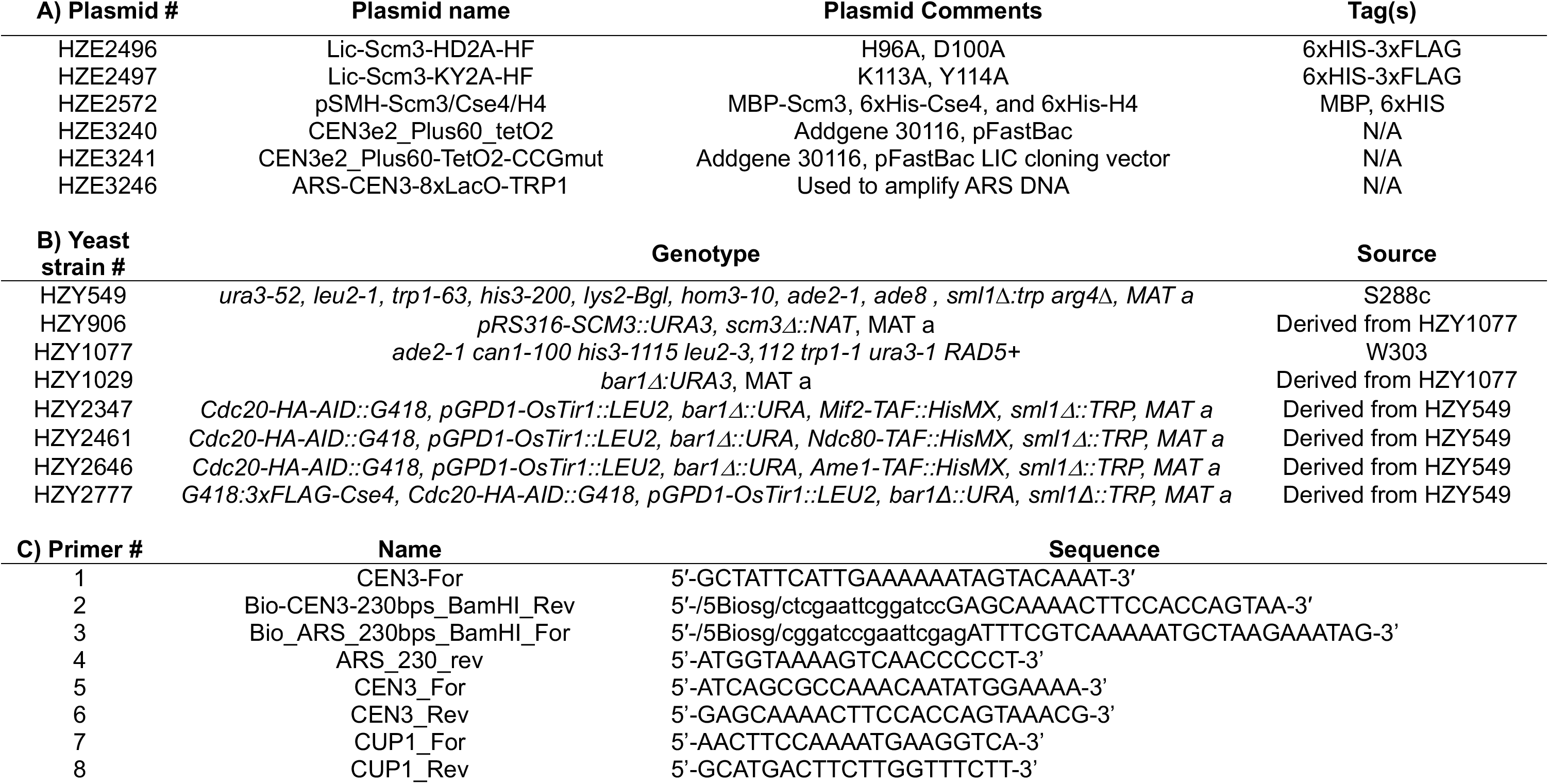
A) Plasmids, B) yeast strains, and C) primers used.

**Supplementary Table 2.** Recombinant proteins used for each experiment.

| Figure | Source of WCE | Recombinant proteins used |  |  | Pre-incubation or purified as complex |
|---|---|---|---|---|---|
|  |  | Protein(s) | Plasmid # | Concentration (nM) |  |
| 1C | HZY1029 | Scm3-6xHIS-3xFLAG, Cse4 has no tag, and 6xHis-tev-H4 | 1792, 1794 | 180 | Purified complex |
|  |  | 6xHis-tev-Cbf1 | 1826 | 635 | N/A |
|  |  | 6xHis-tev-Sgt1 | 1830 | 560 | N/A |
| 1D | HZY1029 | Scm3-6xHIS-3xFLAG, Cse4 has no tag, and 6xHis-tev-H4 | 1792, 1794 | As labeled | Purified complex |
| 1F | HZY1029 | 6xHis-tev-H4, Cse4 has no tag | 1792 | 100 | Pre-incubation |
|  |  | 6xHis-tev-Scm3 | 1754 | 100 | Pre-incubation |
| 2A | HZY1029 | 6xHis-tev-H4, Cse4 has no tag | 1792 | 30 | Pre-incubation |
|  |  | 6xHis-tev-Scm3 | 1754 | 30 | Pre-incubation |
| 2B | HZY2347<br>HZY2461<br>HZY2646 | Scm3-6xHIS-3xFLAG, Cse4 has no tag, and 6xHis-tev-H4 | 1792, 1794 | 100 | Purified complex |
| 3D | N/A | 6xHis-tev-H4, Cse4 has no tag | 1792 | 1,000 | N/A |
|  |  | Scm3-6xHIS-3xFLAG | 1794 | 1,000 |  |
|  |  | Scm3-HD2A-6xHIS-3xFLAG | 2496 | 1,000 |  |
|  |  | Scm3-KY2A-6xHIS-3xFLAG | 2497 | 1,000 |  |
| 3E | HZY1029 | 6xHis-tev-H4, Cse4 has no tag | 1792 | 100 | Pre-incubation |
|  |  | 6xHis-tev-Scm3 | 1754 | 100 | Pre-incubation |
|  |  | 6xHis-tev-Scm3-KY2A | 1758 | 100 | Pre-incubation |
|  |  | 6xHis-tev-Scm3-HD2A | 1762 | 100 | Pre-incubation |
| 4C | N/A | 6xHIS-tev-Ndc10<sup>565-956</sup> | 2463 | 10,000 | N/A |
|  |  | GST-Scm3<sup>1-60</sup> | 2462 | 2,000 |  |
|  |  | GST-scm3<sup>1-60</sup>-5LA | 2469 | 2,000 |  |
|  |  | GST-scm3<sup>1-60</sup>-3LA | 2470 | 2,000 |  |
|  |  | GST-scm3<sup>1-60</sup>-2LA | 2471 | 2,000 |  |
|  |  | GST-scm3<sup>1-60</sup>-2KA | 2472 | 2,000 |  |
| 4D | HZY1029 | 6xHis-tev-H4, Cse4 has no tag | 1792 | 100 | Pre-incubation |
|  |  | 6xHIS-tev-Scm3 | 1754 | 100 | Pre-incubation |
|  |  | 6xHIS-tev-Scm3-5LA | 1778 | 100 | Pre-incubation |
|  |  | 6xHIS-tev-Scm3-3LA | 1779 | 100 | Pre-incubation |
|  |  | 6xHIS-tev-Scm3-2LA | 1780 | 100 | Pre-incubation |
|  |  | 6xHIS-tev-Scm3-2KA | 1781 | 100 | Pre-incubation |
| S2 | HZY2777 | MBP-Scm3, 6xHis-Cse4, and 6xHIS-H4 | 2572 | 100 | Purified complex |
|  |  | 6xHIS-tev-Scm3, 6xHis-tev-H4, and Internal-3xFLAG-Cse4 | 1754, 1791 | 100 | Purified complex |
|  |  | Scm3-6xHIS-3xFLAG, Cse4 has no tag, and 6xHis-tev-H4 | 1792, 1794 | 100 | Purified complex |
| S3 | HZY1029 | 6xHis-tev-H4, Cse4 has no tag | 1792 | As labeled | Pre-incubation |
|  |  | 6xHis-tev-Scm3 | 1754 | As labeled | Pre-incubation |

**Supplementary Table 3.** Antibodies used.

| Antibody | Description | Dilution | Source |
|---|---|---|---|
| α-Ndc10 | Rabbit polyclonal | 1:10,000 | Desai Lab |
| α-Cse4 | Rabbit polyclonal | 1:10,000 | Desai Lab |
| α-Mif2 | Rabbit polyclonal | 1:5,000 | Desai Lab |
| α-Ctf19 | Rabbit polyclonal | 1:10,000 | Desai Lab |
| α-Ndc80 | Rabbit polyclonal | 1:10,000 | Desai Lab |
| α-FLAG M2 | Mouse monoclonal | 1:10,000 | Sigma-Aldrich F3165 |
| α-Protein A | Rabbit polyclonal | 1:10,000 | Sigma-Aldrich P3775 |
| α-Rabbit | Goat polyclonal | 1:10,000 | Millipore 12-348 |
| α-Mouse | Goat polyclonal | 1:10,000 | Millipore 12-349 |

